# Rearranged zebrafish genomic DNA induces zebrafish mutant after microinjection into fertilized egg and preliminary study of the mechanism

**DOI:** 10.1101/460238

**Authors:** Zheming Cao, Weidong Ding, Xuwen Bing, Jun Qiang, Pao Xu

**Author notes:** corresponding author: Key Laboratory of Genetic Breeding of Aquatic animal and Aquaculture Biology, Freshwater Fisheries Research Center,Chinese Academy of Fishery Sciences, Wuxi, 214081, P.R. China, Pao Xu, email-address.

## Abstract

Genomic DNA of zebrafish was first digested incompletely with *Msp I*, and then the fragments were joined to form rearranged genomic DNA. This rearranged genomic DNA was incompletely digested with EcoR I, and the fragments were linked with a long adaptor. Two primers (Gmprimer1 and Gmprimer2) were designed according to the adaptor sequence for two-step amplification. The Gmprimer1-amplified products were microinjected into fertilized zebrafish eggs after purification and a red flesh mutant was observed among 42 surviving zebrafish. We obtained several introduced sequences by two-step amplification. The second set of Gmprimer2-amplified products were purified and microinjected into fertilized zebrafish eggs; all 37 surviving fish were red flesh mutants. We found that the largest amplified band from the mutant from the first microinjection was also present in the amplified pattern from six mutants from the second microinjection. The length of the sequence was 2,565 bp, but it did not encode any proteins. Microinjecting this sequence into fertilized zebrafish eggs produced the red flesh mutant. The sequences differed slightly among different individuals from the second microinjection. Most regions of these sequences were the same, with the exception of a hypervariable region. We mixed 10 such sequences equally and microinjected them into zebrafish zygotes; the findings showed that most zygotes died and the surviving zebrafish were almost all mutants. By genome walking, we found that the site of insertion of the fragment was the same, beginning at position 41,365,003 of the eighth chromosome, and that downstream of the introduced fragment is a conservative sequence of 6,536 bp (named Cao-sequence), starting from a small reverse repeat sequence, not encoding any gene, nor similar to any known regulatory sequence. It has 322 homologous sequences in the zebrafish genome, which are distributed in all chromosomes. We designed two primers within Cao-sequence and several primers specific for different locations upstream of it. Compared with normal zebrafish, we found that the amplified patterns of all mutants in Cao-sequence regions changed to varying degrees. To further understand the effect of the introduced sequence on the zebrafish genomes, we selected six mutants for whole-genome resequencing. The results showed that numerous Cao-sequences from these six mutants were partially deleted and the lengths of the deletions was mostly approximately 6,100 bp, being located at the 5′ end of Cao-sequences. Among them, 43 Cao-sequence loci were commonly deleted from the six mutants (with slightly different locations), and the other deletion sites were not identical. We think that different deletion combinations of Cao-sequence may show different mutation characteristics. The tail part from four red flesh mutants and three individuals of wild type were collected for transcriptome sequencing. TopGO analysis showed that the 4 most significant enrichment nodes were sequence specific DNA binding proteins, sequence specific transcription factors, chromatin proteins and zinc binding proteins. The results of KEGG enrichment analysis showed that the top four affected KEGG-pathways were metabolic pathways, oxidative phosphorylation, citrate cycle and 2-oxocarboxylic acid metabolism.We conclude that deletion of Cao-sequence can affect the expression of a series of transcription regulators and specific DNA binding proteins, then many basic metabolic processes were disturbed which led to mutations.

## Introduction

Our discovery began with construction of a repeatable mutagenesis technique based on the research reports on the introduction of total genomic DNA from distantly related species.It has been widely reported that introducing total genomic DNA from distantly related species into a host cell can lead directly to mutation. Many researchers in China have bred new plant varieties based on this phenomenon. Similar studies have been reported in fish. However, it is unknown how organisms process the introduced DNA fragments and which sequences are integrated into the host genome resulting in mutations. Thus, the repeatability of the mutant traits is low. Some whole-genome sequencing studies have reported difficulty in identifying which sequences can be integrated into the host genome.

Based on the observation that introducing total genomic DNA from distantly related species into a host cell can lead directly to mutation and that most of the distantly related DNA sequences in the host cell are degraded, we developed a new method for no involving the introduction of distantly related DNA. Specifically, we directly rearranged the genomic DNA of common carp into new fragments with a different arrangement and introduced them into fertilized carp eggs. The results showed that these fragments can lead to common carp mutants. However, this method suffered from many problems and we did not determine whether this phenomenon can be repeated in other fish. Thus, the objective of this study was to further improve this method by applying it to zebrafish.

## Results

A total of 200 fertilized zebrafish eggs survived after microinjection of the rearranged DNA fragments and 32 individuals were obtained after 3 months of feeding. A zebrafish mutant with flesh that was red in color was detected. As shown in Fig. 1, the zebrafish with red flesh is the mutant and the blue-gray one is a typical zebrafish.

**Fig. 1.**
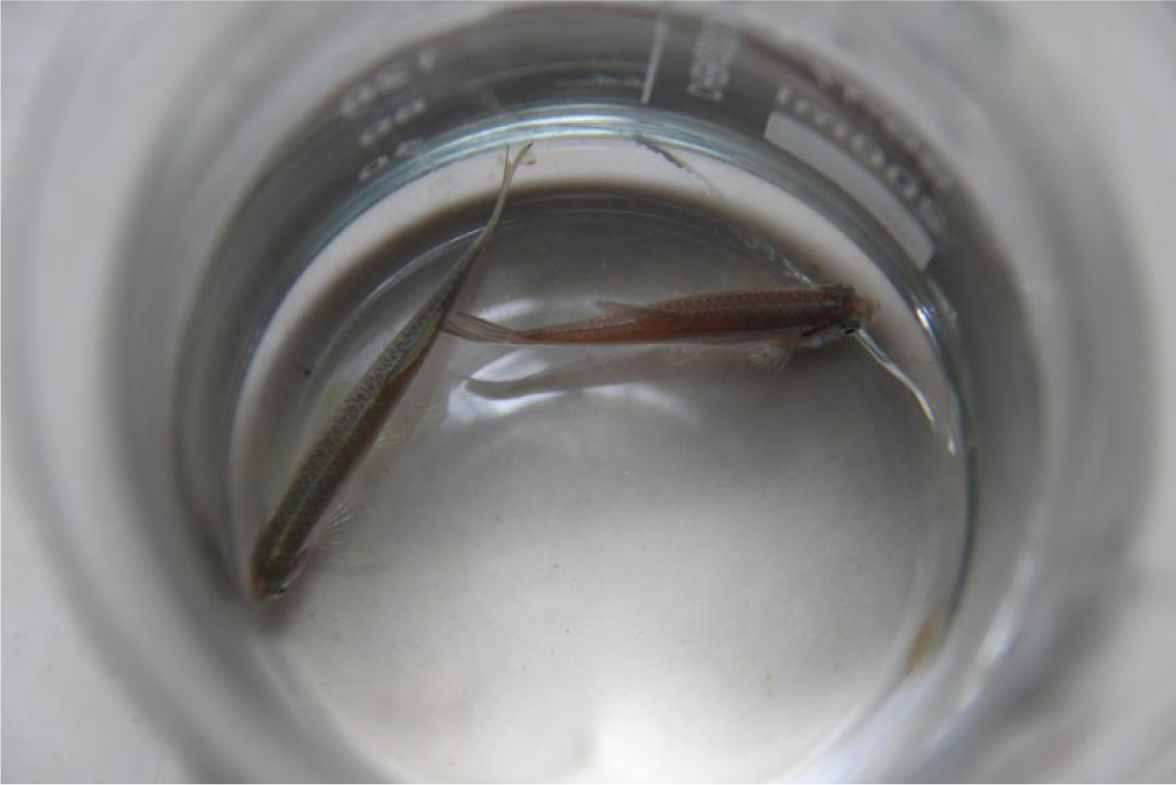
Mutant (red) and typical zebrafish

We obtained several sequences after the two-step amplification. The largest amplified band from the mutant appeared in most mutants obtained after the second microinjection of purified PCR product from the second amplification, which may be related to the body color mutation (Fig. 2). The control shown in Fig. 2 is not the mutant with red flesh after microinjecting the rearranged DNA fragments.

**Fig. 2.**
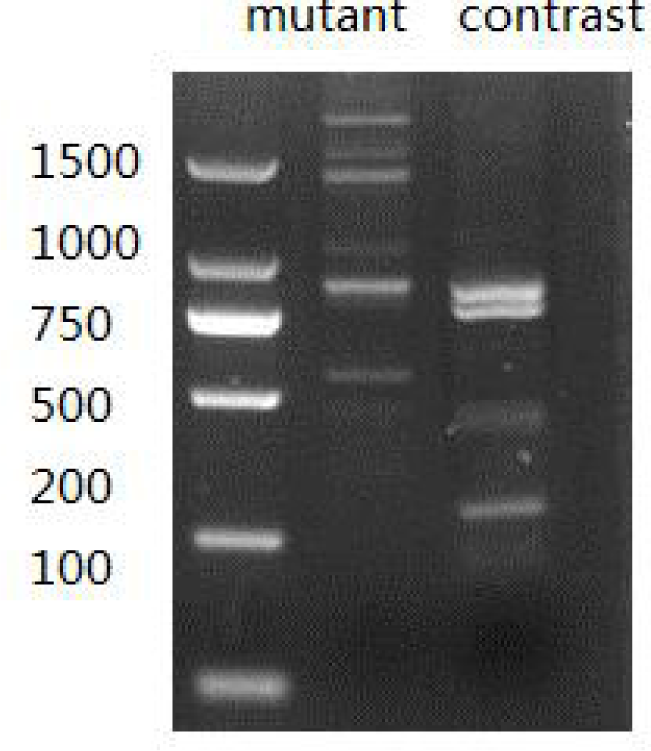
Second amplification result

A total of 200 fertilized zebrafish eggs survived after microinjecting the amplified sequences, and 37 fish were obtained after 3 months of feeding. All fish were red flesh mutants.

We extracted the genomic DNA from six mutants and found that five of them had a common amplified band with different amplification efficiencies (Fig. 3). We hypothesized that this might be related to the body color mutation.

**Fig. 3.**
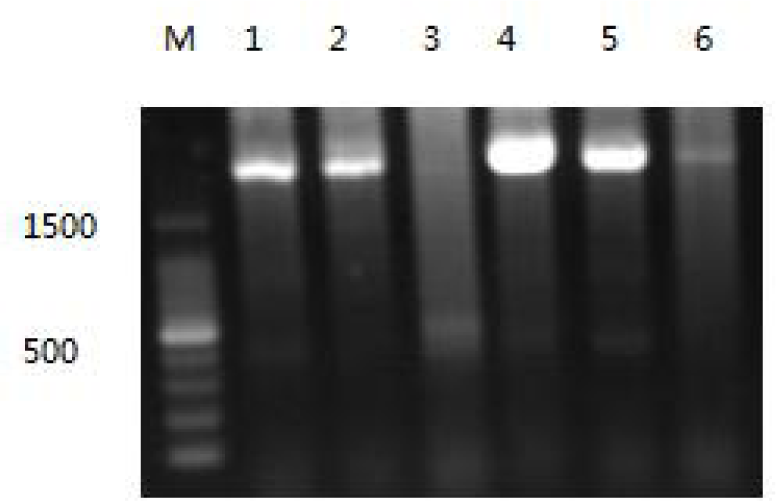
Result of amplifying introduced sequences from six red flesh mutants

We purified the largest amplified band shown in Fig. 2. The result of sequencing the common amplified band was as follows:

ACAGGAAGCCGAATTCATGCAGTTTGTCTTCATTCAGAGTAAACGTCATCCATGGCGAGAGAATCCAGAGAGAGAATAATGACCCGTGTTGTGACAGAAAAAGCACAGCATATTTAGACTGTGATGTAAAGTCACAAGGGCTGACCTGTTGCATTCATACTCTCAAACTGAGAGGAAACAGAAACGACAAGAAAGGTTTGTATACAGGAAGCTTGGGGCATATATCTTATTATGCTGAACCACTAGAGGCACATTTGCCACAAAGTAGCATTACACCACTTAATAAATCAGTGTTCACAGCGGTATGAAGCACTTCCCATATAGTATGTGGCAAAATGTTTATACAGAGGT CAGACAAGTCCCACGTATTAATTTATTGATTTATTGGGGGAGAAA **GTTTTAATGTATTA CTTGTTTAATGTGAGGTAAGGGTTGTGATCATGACATTAGGATTCTCCCAAATTCTTTAAACATGTTAGCCTTCAGAGCTGCTTTAAAGATGCCACAGCGTTCATTGACTCAATAAGTTATATTCAGAGTTGTGTTTTGGCTTAAATACAGATTCAAGAGTTCACACTTAGCTGATGATTGGTAATAAGCTAGTTTGGCATGCTCCCTCCCGTTGCATGGCGAGAGGAGAGTTTGAGCTCGGGTAGATCTCGAGAACCCCCCCAACACCAACCGCTGCGTGAGGGTTGCTAAGGTGATGCGTTGCTGACAGAATTTAATTGAATTTTGTGCTAATTTGGTTTAGTCAATTTACTTAAGTTCATGTTTTTGGGAGTGTGGGAGGAAACCGGAGGACCCGGGGAAAACCCACGCAAGCATGGGGAGAACATGAACTCCACACGAAAATGCTGTCCGTGTGGGTAGATGCCCGAACCGGAGGCATTTTTGCTGTGAGGCCGCATTGCTAGCCACTGGGCCACCGTGCTGCCCTGTAGAAAAGAGGAGGAGATGGGGTGGAAGGGGGGATTTTTCAAGACAAAGATGACTAAAGGTAAGGTGTTTTATACTTGGTTAGGAATAGTCTGATTGGCGGTTCGTACATTAGCTTGACTCGGGGCCAGCCGTGTCAATCATGAACACGTGCTCCTCTCGAAATTAGTTTATGAATAAACTTTACTAAATTTAAGTTACAGTCGACAAAAGGAACCAACAAAGATGTATCTAGAGTTGTATTTTGGAGCTTTAACATACAGGTGTTGGAGAATAAAACTGAAATGCCTGGTTTTAGACCACAAATATGTTGTAGGGCCTCCTTTTGCGGCCAATACAGCATCGCTTTGTCTTGGGAGTGACACAAGTCCTGCTTAGTGGACAAAGGGATTTTGAGCCTTTCTTTTTGCAGAATAGTGGCCAGGTCACTGTGTGATGCTGGTGGAGAAAAACAAGGCCTGACTTGCTCCAAAACATCCCAAAGTGGCTCAATAACATTTAGATTTGATGACTGTGCAGGCCATGGGAGATGTTCAGCTTCACTTTCATGTTTATCAAACCCCTCTGTCACCAGTCTTGCTGTGTGTATTGGTGTATTATCATCATGTTCAGGAAACAGTGTTTGAACCATTAGGTTCACATCGACCTCCAGAATTGTTTGGTCACAAGCTCAAATATTGTGTTTAGGGAATGCCTTGATATTGCAGCCCAAACCATCACTGATCCATGCTTCACTTTGGGCATAAACAGTCTGGGTGATACACATCTTTTGGGTTTCTCCACACCGTAACT CTCCCAGATTTGGAAAAGACAGTGGAGGTGGACTCATCAGAGAACACTACATGTTTCATATTTTGTCTGCAGACCAACATTTTCACTCCTGGCACCATTGAAACCAACGTTTGGCAAGGTTTGTTCAAGGTTTGTCATTGTCCAACCCCTGTGCATGAACACACAGGCAGGAATTAATAACAATCTTTGCATAAACAGATGAGTTTACTGGCATTCGTTCAC TCAAGAAATAT** CACAGAAAGAGTAATGTTTCATGACGCATAGATGTGCA **TTAAATAGC ATTTTGCATGTGTTTGTTGCCTCAATACAAACATGCACTGAAGATTTTACAATATGCATAGTATACTAACACAAGATACATTTGAATATGATACAGCTGAATAAATTTGTAATCGGTATATCTCGCTTTTTGGTTTTGGATAAGTTATGGGTGCATGGGTGCTTGCCTGCTTCACAAAACTGCTCCTTTTAGTCTACCTGTCTCTTACATTTCAAAATAAGAATCCCACTTACATGGTTTGTTGAATTCAAGCTTAGAAAAATGTAAAAATAAGAAGAGAATATAATGTTGTAATCTCACTCGGAAATGTGACCAATCAAACACAGTCTGGAGCATGCATATTAATAATGCTGTGTAGATCTTAAGAGTTGGTAGAGGTGGCTCCCTTTAGCATGTTTTTTTCACCAGTGGTTTTCACTCAAATTATTTACATGAGAACGAGGAAACAAATGGGGAACTTGTTTTCAGCTATGCCTGGGTATATCCAGATTTTCATTCCATGGCACTTTAAGGTTAAGTAAGGTCAAGCTTTATTGTCATTCGGCTACATGTGT**

The sequence length was 2,565 bp and it was divided into four segments. The first (396 bp) and second segments (1,568 bp) had homologous sequences on the eighth chromosome (41,365,386-41,365,781 and 41,368,513-41,370,080), the third segment was a short sequence (38 bp), and the fourth segment (563 bp) had a homologous sequence on the eighth chromosome (41,376,421-41,376,983). They were not all coding sequences and none was related to any known regulatory sequences. The sequence was made up of four fragments produced by restriction endonuclease *MspI*, but we did not find any typical MspI cleavage sites at the junction of these four parts.

We selected 10 individuals from 37 mutants after the second microinjection for genomic DNA extraction and obtained 10 sequences amplified by primer Gmprimer2 from 10 genomic DNA samples. We found that the sequences obtained from different individuals were slightly different, with the differences being concentrated in a hypervariable region, as shown in Fig. 4.

**Fig. 4.**
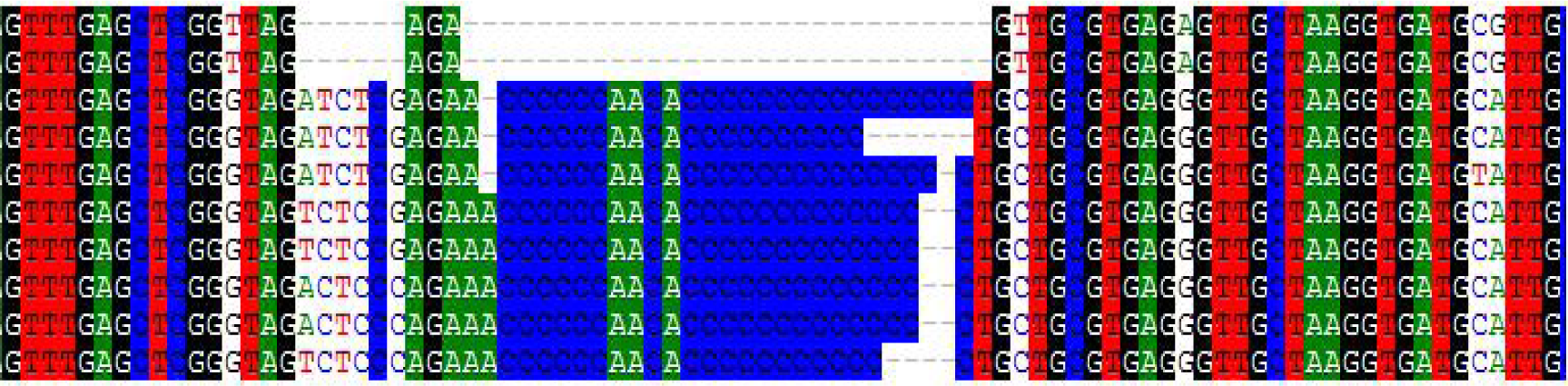
Comparison of 10 introduced sequences amplified from 10 mutants

We purified the 10 amplified bands and mixed them equally, and then microinjected them into fertilized zebrafish eggs. The results of microinjection are not shown in Table 3.

**Table 3.**
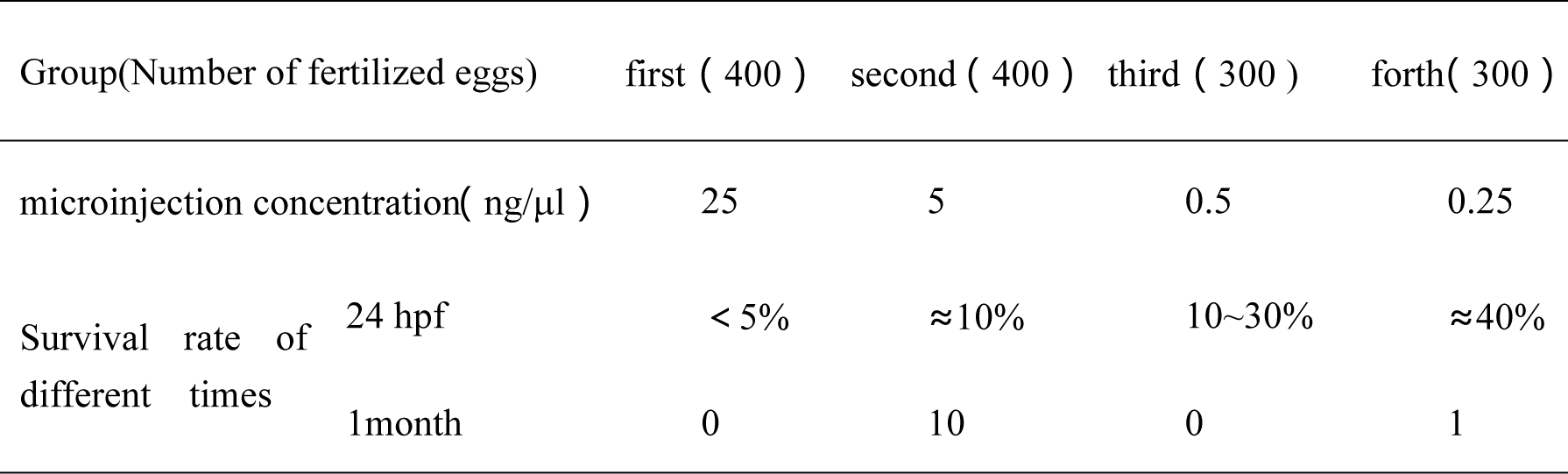
relationship between microinjection concentration and survival rate

Most fertilized zebrafish eggs were killed after injection at 24 hpf (24 hours after fertilization) in all groups and the survival rate was less than 1% when the fry were raised to 1 month old. The maximum difference in injection dose was 100 times. The survival rate at 24 hpf increased significantly when the injection concentration was decreased, but the survival rate at 1 month old was basically unchanged.

Among the surviving zebrafish, numerous mutants were found, two of which are shown in Fig. 5 and 6. A mutant that could not grow up and a mutant with curvature of the spine was also observed. In the remaining individuals there were no marked changes in appearance, but after months of feeding, three emaciated fish that were unable to eat anything and subsequently died were identified. It is possible that these mutants showed signs of mutation after maturing.

**Fig. 5.**
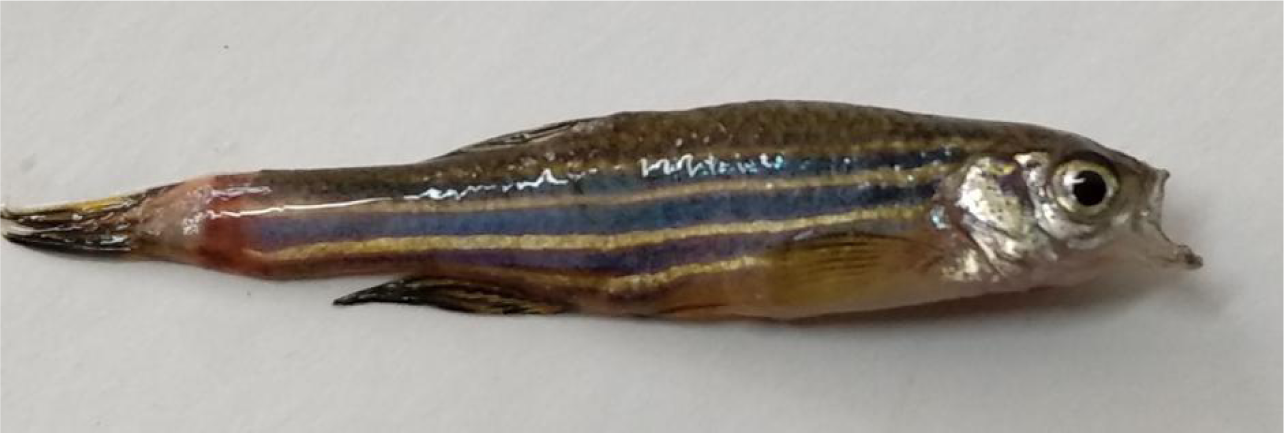
Mutant unable to close its mouth

**Fig. 6.**
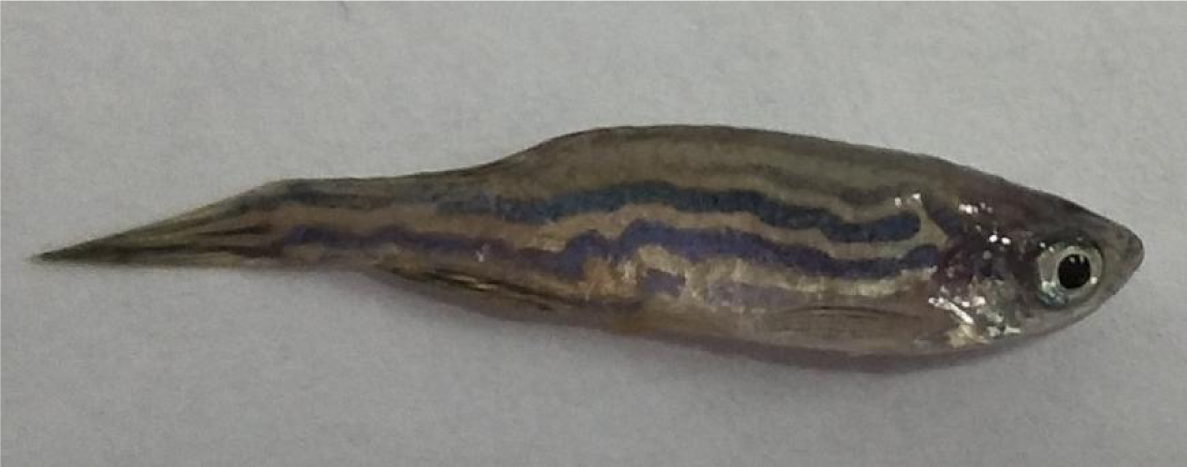
Mutant with curved stripes

We obtained the sequences upstream and downstream of the insertion site by chromosome walking. We found that the region upstream of the insertion site began at position 41,365,003 of the eighth chromosome, only 383 bp away from the location of the introduced sequence. We repeated this analysis in 10 individuals and obtained the same results. It was thus concluded that the first part of the introduced fragment basically determines the position of the insertion site. The downstream sequence started from a conservative sequence. The conservative sequence is as follows:

AACGTTCGAGGTTACAGAAGTAACCCTTCGTTCCCCGAGGAGGGGAACGGAAGTGCCATGAATGGGAGGATTCGGATCAGAAGCCGCTTATCTGGAGAGTATTGAACGGGCCAATGAATGAAATTAATTGGCAGCGTAAGCTTGCGCAGGTGTGCGACATCTGCAATTATCTCAGCATATAAGCACACCTGAAGCCAGCAGACGCCATCCTTTTAAGCTGAAGAGACTTTCAAACAGCTAAGGGACAGTCATTATGGCGACGGAATATGGCACTTCCGTTCCCCTCCTCGGGGAACGAAGGGTTACTTCTGTAACCTCGAACGTTCCCCTTCGGTTGGGGAACTTCAGTGCCATGAATGGGAGAATATGGAAAGCGCCGTAATGACTGCACCTTACCAACACCCCCGATGAGGAGATAGTCAAGCAAGCGTGACGCACCCACATCATGGGGGGCGCGGTCCTCCAACGTGTCCCTGGCCCTAATTTATCCTACTTCAACAGAAGTTTTACGGATTTAGATATATTTTTTGGGAAGTCGTGAGCATCTAGATAATTCTAGGAAATACGACAGTACGTTGGGAAGCGTGCAATCCCGATAGGGAGGACGCTGCGGAGGCCATCCGTTACCCAAGGGGGGGATAGATGGCAGAATTTACCTATGGACTAGCCCTAAAAAGGGGGAGTACGCATAGCAAAAGAGTGGTTAGCGGAGAGGGAAGACACGGGTCCGCCCAGGGGGGGGACTTAACCGTGGCGGAATAAGCATATGGGATCGCCTAGTGGGGATCACGCATAGCAGGCACCTATACCCAAAACGCGGGCTGACCAGCGGGCAGACCTACAACGTAGTGGGCCAGCAAGTGACTCCTCCGCTGAGTCAGTGCTGGGGGCCACGGAGGAATCTGCAGGGCTCACCTGACGGGGAACTTTACTGACAGATAAAAAAGGCGCACGTACCTCCGTGTTAGGGAGAATGGCGCCGCAAGCGTGTTTCAACACCCTACCGAGTTGTCTCCTCAATCACCAAAGGGTTACCTAATACCCTTGAGGAAACCGGCTCCACTCGCAGATTGTAAAACCTTGCAAATGTGTTGGGTGTCGCCCAGCCCGCAGCTCTACAGATGTCTGTTAGAGAGGCGCCGCGTGCACGCGCCCAAGAGGATGCAACGCTCCGAGTGGAGTGTGCACGAACTCCCGGGGGACACGGCTGACCTCGACTCGAATAAGCGAGTGAAATGGCATCCACAATCCAGTGGGATAATCTTTGTTTTGATACGGCACTTCCCTGCTGCCGACCGCCATAACAGACAAAGAGCTGCTCAGATGATCTAAAATTCTGAGTACGGTCCACATAAATGCGCAGAGCGCGAACTGGACAAATTAAAGAAAGGGCTGGGTCTGCATCCTCCGGGGGCAGCGCTTGCAGGTTCACTACCTGATCTCTAAAGGGGGTGGTAGGAACCTTGGGCACATAACCGGGGCGGGGTCTCAGGATGACGTGAGAGTAATCCGGCCCGAATTCCAGGCACGAGTCACTGACCGAAAATGCCTCCAGGTCCCCGACCCTCTTGATGGAGGCCAACGCAACCAGCAGAGCTGTCTTCAGGGACAGAAATCTTAGAGATACTGATTCGAGTGGCTCAAAGGGATCGGATCGCAGACTCGTGAGAACGAGGGCGAGATCCCAAGAGGGCATGAGAGGGGGGCGAGATGGATTAATTCGCCTAGCACCCCTAAGGAACTGGATGACCAGGTTATGCTTTCCCACGGTGCCGCCAGCTACCGCGCTATGATAAGCGGAGATGGCGGCCACGTAAACCTTGAGAGTGGAGGGCGACAGCCTGCTGTCCAACTTCTCTTGAAGGAAAGAGAGCACAACACTAATCTGGCAATTTCGGGGGTCTTCTCTGCGAGAGACGCACCATTCAGTGAATAGACTCCACTTCAGGGCGTAGGCGCGCCTCGTGGAGGGGGCTCTAGCCTGAGTGATGGTATTAACCACCGCAGTCGGTAGGTTACCTAAGTCTTCCTCGCGTCTAGGGACCACACGTGGAGGTTCCAAAGATCGGGGCGAGGGTGCCAGATGGTGCCCTGTCCCTGAGAGAGTAGGTCCTCTCTCAAAGGGATCCGCCAGGGGAGGGCCGTCGCGAGGAGTGAGAGCTCTGATATCCAGGTCCGGTTGGGCCAAAGGGGCGCAACTAGCAGAACCTGTTCCTCGTCCTCCCTGACCTTGCACAGAAACTGCGCGAGCAGGCTCACTGGGGGAAACGCATACTTGCGCATGCCCCGAGGCCAGCTGTGGGCCAGTGCATCCGTGCCGAGAGAGCCCTCGGTCAGGGAAAAAAACAACTGGCAGTGAGCGTTCTCGGGGGAAGCAAACAGATCGATCTGGGCCTCCCCGAATCGCGCCCATATCAGCTGAACAGACTCGGGGTGGAGTCTCCATTCTCCAGGGCGTAACAGCTGTCGTAAGAGCGCATCGGCTGCACGATTGAGCGTGCCTGGGACGTGAATGGCGCGCAGCGATTTCAGCCGCGGGTGACTCCAGAGGAGCAGACGGCGGGCGAGCTGAGACATGCGGCGAGAGCGCATACCCCCCATGCGGTTGATATACGCCGCCGCCGCCATACTGTCCATCCTGACCAGCACGTGTTGCCGCTCCAGCACCGGTAAAAAATGGTGGAGAGCGAGGAACACTGCCAACAGCTTTAGGCGATTGATATGCCAATGCAGCTGGGCACCCTTCCAGAGGCCCGCAGCCGCATGCCCGCGACACACGGCCCCCCAACCCGTGTTGGAAGCGTCTGTTGAAACAACAACATGGCTGGACGCCTGTCCTAGAGGCACACCGGCCTGTAGGAACGAGGGGTCGTTCCAAGGGCTGAGGGCGCGGCGACACAGCGCAGTAACCGAGACCCGGTGTGTGCCCGCGTGCCATGCGCGTCTGGGGACCCGATCGTGAAGCCAGTGCT GAAGTGGTCTCATATGGAGCAACCCGAGCGGCGTGACGGCGGCTGCGGATGCCATATGCCCCAGGAGCCTCTGAAAGAACTTCAGTGGGACCACTAGTTTGCTGTCGAGCTCCCTCAGACAGTTCAGCAACAGGCGAGCGCGTTCCTCGGAGAGGTGCGCTACCATGGTGATCGAGTCCAGCTCCATCCCGAGAAAAGAAATCCTCTGCACGGGGGCGAGTTTGCTCTTTTCTCGGTTGACCTGAAGCCCCAGTAGGCGGAGATGCCGAAGCACCTTGTCCCTGTGCATAATCAATTGCTCCCGCGAGTGGGCTAAAATCAGCCAGTCGTCGAGATAATTGAGTATGCGAATGCCCGCGAGCCGAAGGGGCGCTAGGGCACCCTCCGCGAGTTTGGTGAAGACCCGCGGAGACAGAGAGAGCCCGAAGGGGAGGACCTTGTACTGCCACGCTCGACCCTCGAACGCAAACCGCAGAAATTGGCGGTGGCGTGGAAGAATGGAGACATGGAAATACGCGTCCTTCAGGTCTATGGCTGCAAACCAATCCCGAGGACGAACGCATTGGAGAATGCGCCTCTGCGTGAGCATTCTGAACGGCAGCTTGTGCAGACAGCGGTTCAAAACGCGCAGATCTAGGATTGGCCGTGACCCACCGCTCTTTTTGGGTACGATGAAGTATGGGCTGTAAAACCCACTCTCCATCTCGGCTGGAGGAACCGGCTCGATTGCACCCTTCGCCAGGAGGGCAGCAATCTCCTCTCGCAAGACAGGGGCGGACAGGGGGTTGACCCTGGAGAAATACACGCCCGTAAACTTGGGGGGCCGTTTCGCGAACTGAATCGCGTAACCGAGTCTGATTGTGCGTATGAGCCACCGCGAGGGGCTGGCCCGCGCTAACCAGGCAGGCAGAGCCCTCGCTAATGGAGTCATCGCTACAATCGCTGACGTACCAGCGGTGGGGCAGCGCGGAGTGGGTGTGCTGATCCGGGAAGCACGAGGGTCCCGCGGAAGAGCTGGAGGAGAGCGATTCGCTCTGGCTCCGCACCCTGACTCCGGAGAGGGGGCTGGAGGGCTGAGGAAAGGGAGACCGTCCCCCCAGCGCTCGTGAGCCACTGGCGAAGCACGTAGAGTCTGCGAGCCGGAAGGCAGCGCGTCCCGGACTGGCAGAACTCGTGCAGTCACATCCGGGGAAGGGAAAAAGTGCTCTTTTACTGATGTTTTGGTGGCACTGAAAAGTACCGTTGTTATCGGGGCCCCGCCCTCCAGCGGGGAAAGAGCAAGTTTCCTCTTCTCCGGATGGCCTGTCTCAGGGACGCTTCGCGGTCCGTTTACCGGACTTAACGGCGCCCTGGGCAGGAGGGGCTGCCTGCTTTCGAGGTGAACGCCGCGCCCGCTTGGCCGGAGGCGCAGGCGGGGGCGGGGCAGCACTCGTTGGCGGGCGCCCTCGACGAGGAACAGCGGAGGTGGATGGCTCAGCGGGCGGAGCGGGCTTACGGCCACGCCGATAGATGACATTGCCCATCGCATCCGACTGCTCTTTCACCGCCTTGAATTCCTGGGTGAATTCACCGACGGTGTCGCCGAACAGGCCAGCCTGGGATATGGGCGAGTCAAGAAAGCGAACTTTGTCAACGTCGCGCATATCAGCCAGGTTTAGCCAGAGGTGGCGTTCCTGAACCACAAGTGTGGACATCGTCCTCCCCAGCGCACACGCGGCGGACTTAGTAGTCCGAAGAGCATAGTCGGTCGCGGTGCGCAGCTCATGTAATAAGCTTGGGTTGGACCCACCCTCGTGCAGCTCGGCCAGCGCCTGCGCTTGGTAGCGCTGGTAGGTGGCCATCGCGTGCAAAGCAGAAGCAGCCTGGCCCGCAGCCTTATAAGCTCTGGCTCCGAGGGAGGCAGACAACCTACAGGCTTTGGACGGGAGGCGGGGCAAACCCCGCCACGTAGAGGCGCCGCGCGGACAAAGATTGACCGCGATAGCGCGCTCCACTGACGGGATCGCCTCATACCCCCTGGCAGCTCCGCCGTCAAGGGCGGTGAGGGCGGAGGCACTCGCAGCACGGGCAGAGAAAGGTGCCCTCCAGGACTGCGTGAGCCTACTGTGCACTTCCGGGAAGAAGGGGACGAGAGGCTTCGAAGGCTTCGCCTTCTGGTCCTCTACGTAGCACCCATCTAGTCGGTCCGGCCGCGGAGCTGGGGGATAAACCATCTCCAACCCCCAGGCCGAAGCAGCCCGGGAAAGCACGGCTAACATGTCCGCTTCAGGATCCGATTTGACAGCGCTCACCTGCCCGGAGGGGGCGAGCGGGTACGGATCTTCGTCGGACAGTGAAAGCCCACCCTCCGATGCCGCGGAGGACATCTGATCTCCGGTGTCAGCGAGTGTGAGAGCTACCATCTGGTTAGACGGATCACTCTTACCACCCGAAGCTTGGATGGAGCGCCGTGAGGAGCGAGAGGTCCGCGAGCCCGTGGGCGGCGGATTAGCTCCCGCTGAAACCCTCAGATCTGCCCGAGCGCCCGCTGCTTTTTTAGAGCAGGAGGCAACTGGGGTGGCTCGCTCTCTTGCGAAAGTTAGCCGCGATCTTAGCTGTGCAACGGTCATGGCATCGCAATGACGACATGAACCGCCCGCGAGCACCGCATTAACATGCTGGACCCCCAAACATGCAATGCAGTGATCGTGTCCATCATCCGGAGACAGGAAACCCATGCATCCAGAAACGCACAGTCGGAGCGCCATCCTGAAAAGGACGTGCTGCACGACTGTGTTGCTCTTTTAGGAAAGTTGCAACTATACGCACCGCTCTGGAGGACCGGACCCAAAGAACCGCAGGCAAGGGAGTAACCCAGCTCGACCGTCTGCCACCGCGAAGACCCACTCTGGACCGGGAGACACACTCGTTCGCTCTGAAGTGCTGATCAGCAAGAGGAACCCTCATCGACTCACTCAGAAAGGATCTGAAGCGAAAAGGATGGCGTCTGCTGGCTTCAGGTGTGCTTATATGCTGAGATAATTGCAGATGTCGCACACCTGCGCAAGCTTACGCTGCCAATTAATTTCATTCATTGGCCCGTTCAATACTCTCCAGATAAGCGGC TTCTGATCCGAATCCTCCCATTCATGGCACTGAAGTTCCCCAACCGAAGGGGAACCTATAATTAAGAGGTCCAAGAAACATGTGCAGGTTTTGATGAATAGGCAACTTTATGATAATACTTTACATTTAAGTACAGCAGTTTTCTTTTAACTTTTTAAGATCAGTGCCAATTTAAATTAACTTTTAAAATCACATTTTAATTTAAACATTGACATTTTATTTATCTATTTTTTTTACCTGTAAGTGTGTTATTGTTCAAACTATTTAATATGTGGCTGACAATAATAAATATTCATAATAATAAGAATTAGTCTGTAGTTTTGTAGTAAGCTGTAGCTGCTTAGTTGGGCCTACTATGTATTTCAATACCAGTCATTATGGTGGTACTTGGAAAGGCATTTTTTTCTGAGGTGGTACTTGGTGAAAAACGTTTGAGAACCACTGTTCTAGAGGAACAAG

This sequence does not encode any protein and is not similar to any known regulatory sequence. We named it Cao-sequence. A total of 322 Cao-sequences were found in the zebrafish genome. As shown in Fig. 7, Cao-sequences are widely distributed in all 24 of the zebrafish chromosomes. They are most commonly distributed on chromosome 7 (21 Cao-sequences), and most rarely on chromosomes 10, 15, 22, and 25 (7 Cao-sequences each).

**Fig. 7.**
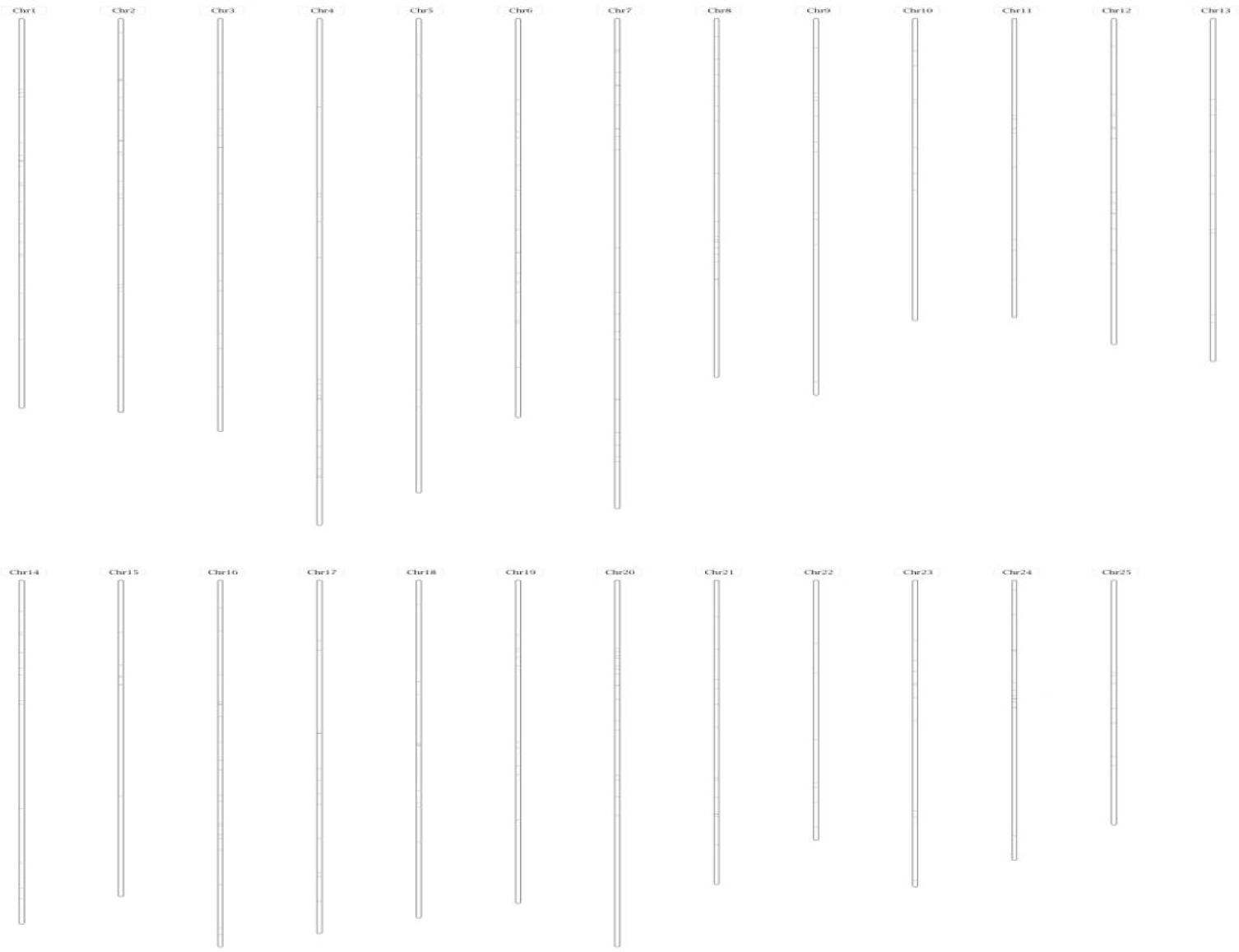
The distribution of Cao-sequences in the zebrafish genome

To study the effect of the introduced sequence on Cao-sequencess, we selected 11 Cao-sequences in the 8th, 9th, 10th, 11th, and 12th chromosomes and amplified the sequence upstream of them to investigate the differences in amplification. Because three parts of the introduced sequence were located in the 8th chromosome, six primers were designed for the 8th chromosome, while the rest were for the 9th, 10th, 11th, and 12th chromosomes. The amplified results are as follows.

**Fig. 8.**
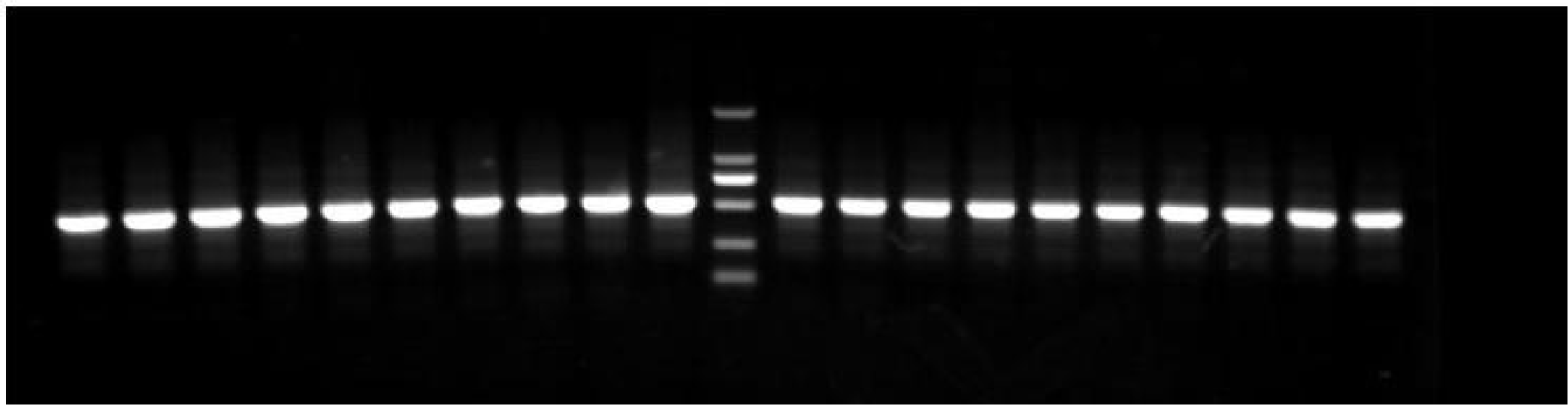
Amplification pattern of primer 8-364

**Fig. 9.**
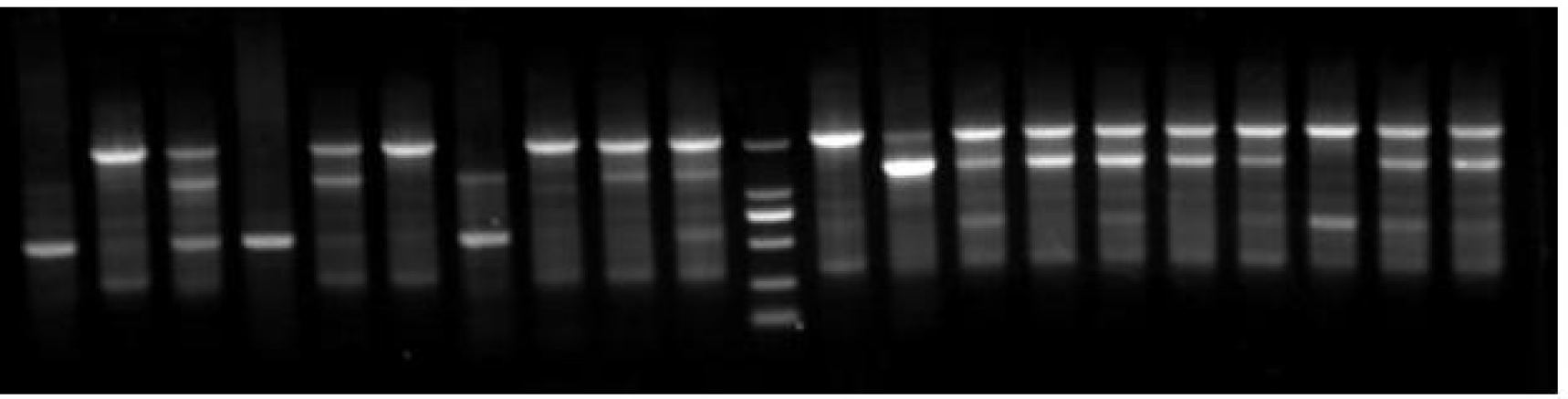
Amplification pattern of primer 8-337

**Fig. 10.**
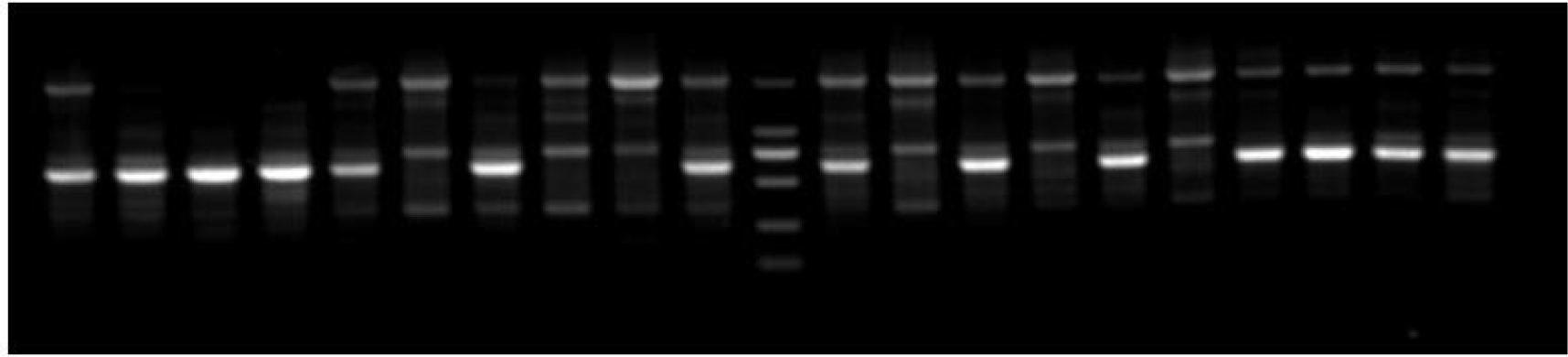
Amplification pattern of primer 8-307

**Fig. 11.**
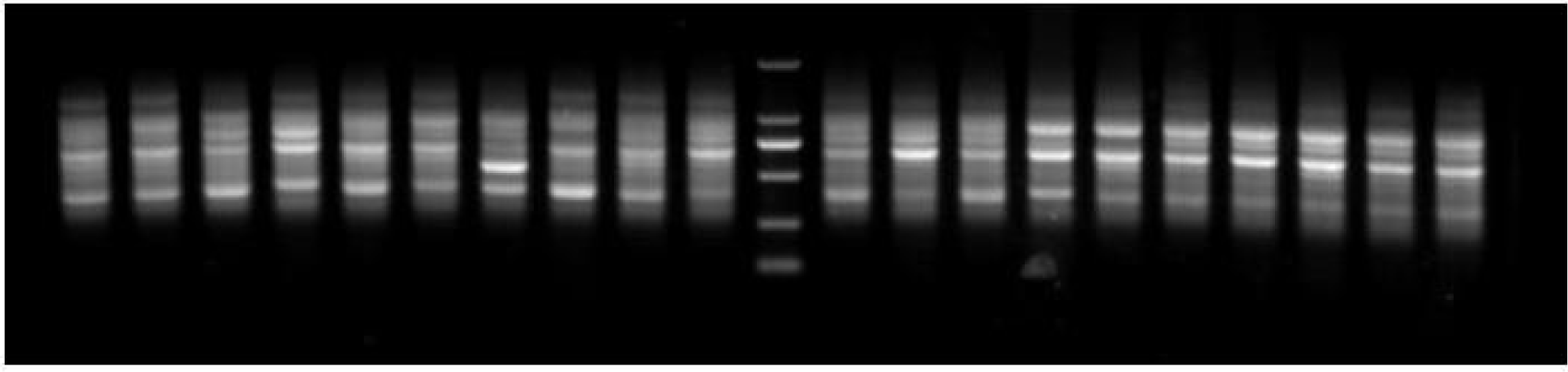
Amplification pattern of primer 8-154

**Fig. 12.**
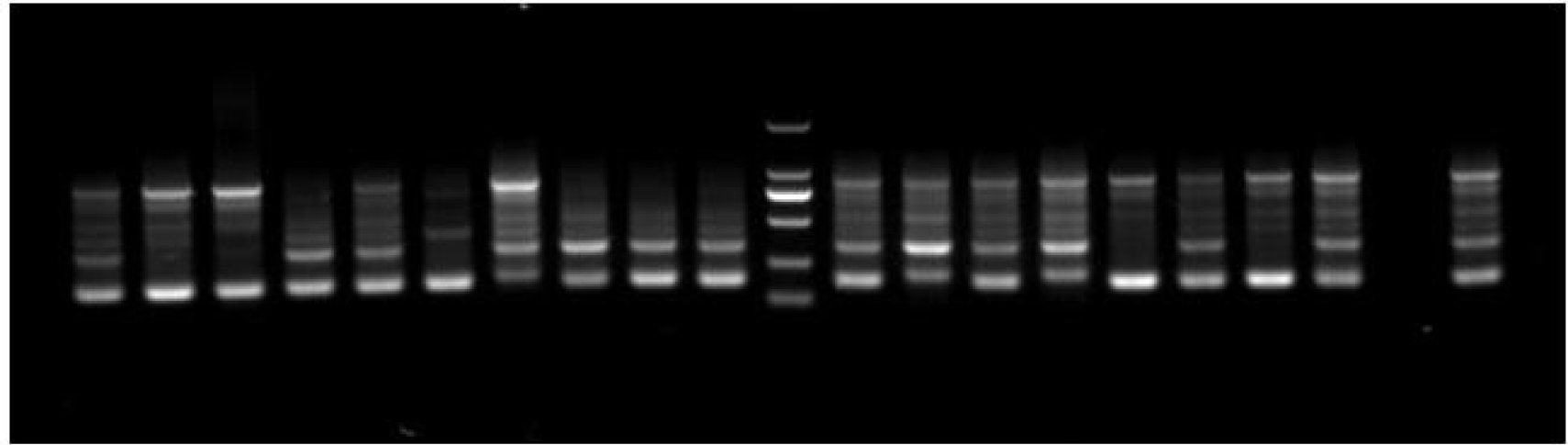
Amplification pattern of primer 8-838

**Fig. 13.**
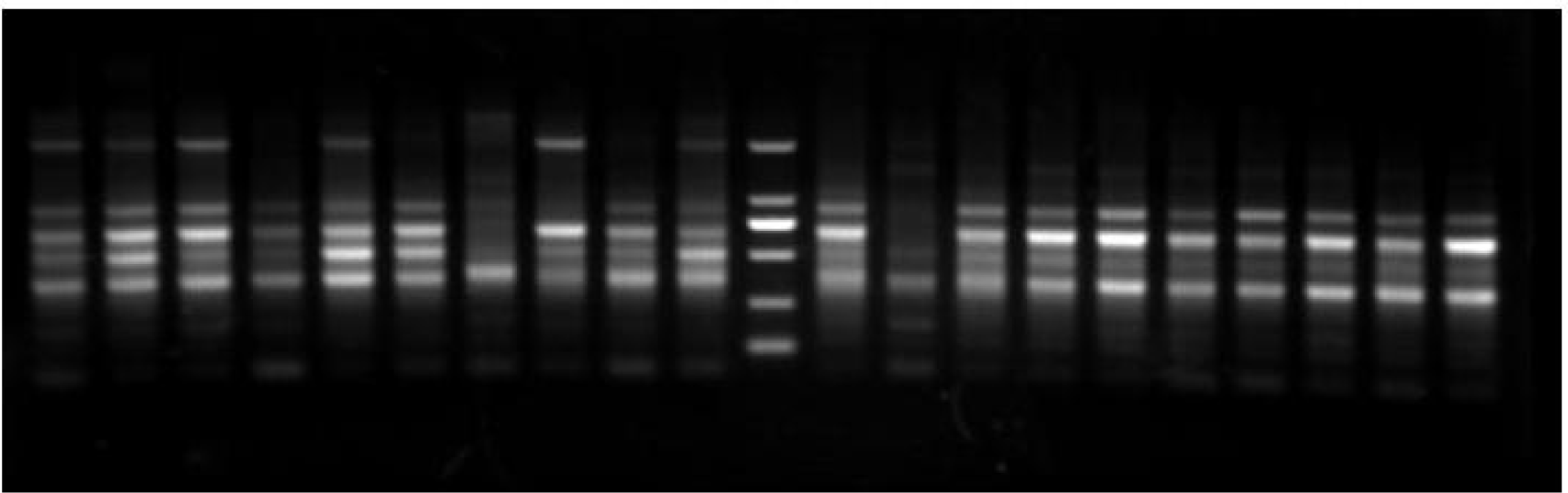
Amplification pattern of primer 8-611

These six electrophoretic maps amplified by primer 8-364,8-337,8-307, 8-154, 8-838 8-611show the results of amplification of sequences upstream of six Cao-sequences from chromosome 8, along with 10 control individuals on the right. We found that the amplification patterns changed in the 10 mutants, except for with primer 8-364.

**Fig. 14.**
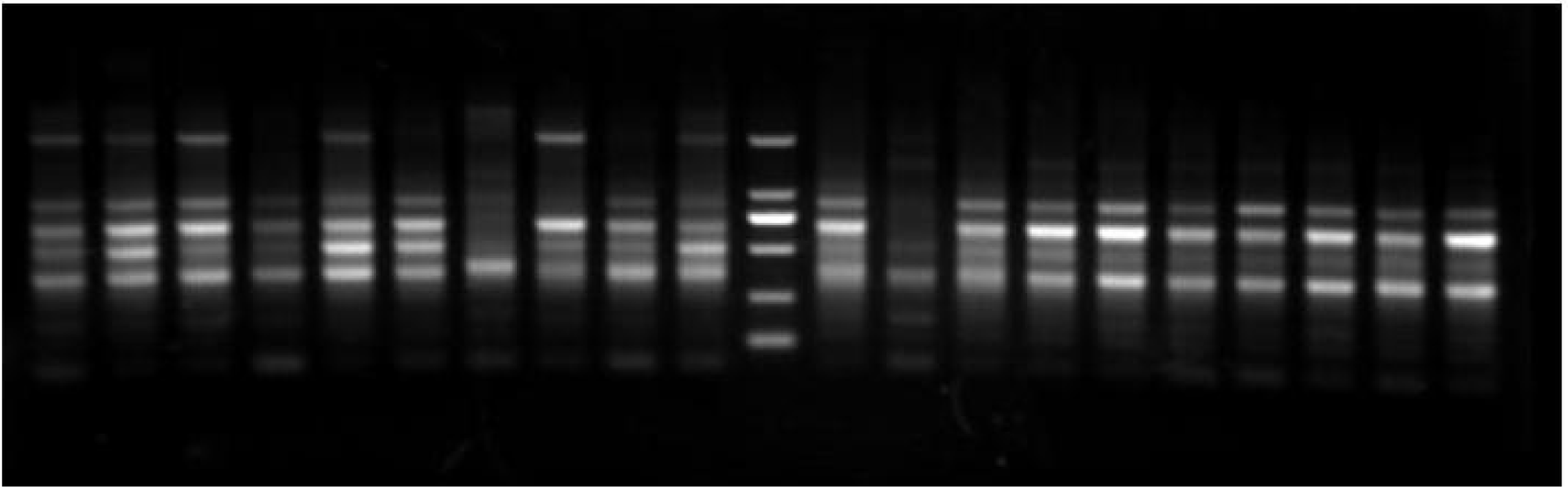
Amplification pattern of primer 9

**Fig. 15.**
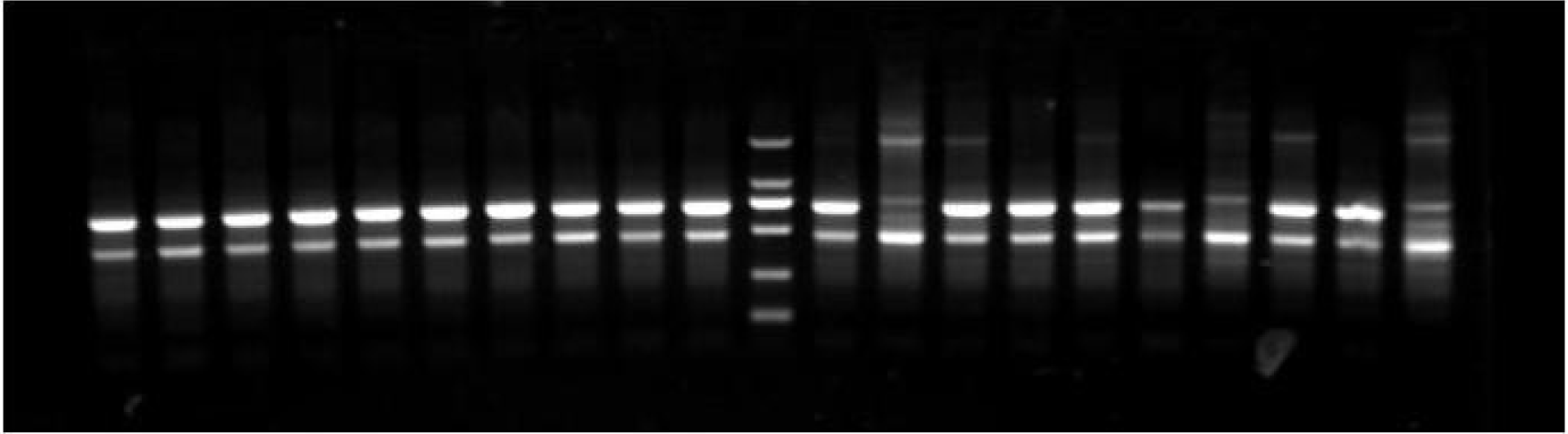
Amplification pattern of primer 10

**Fig. 16.**
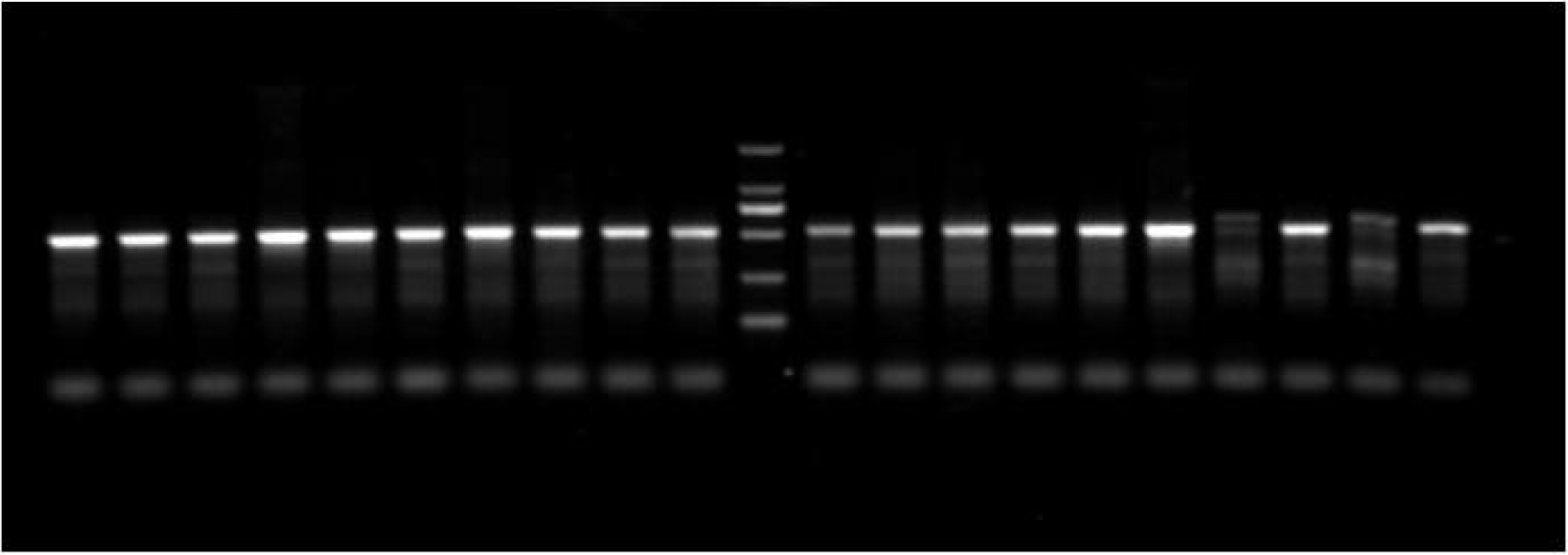
Amplification pattern of primer 11

**Fig. 17.**
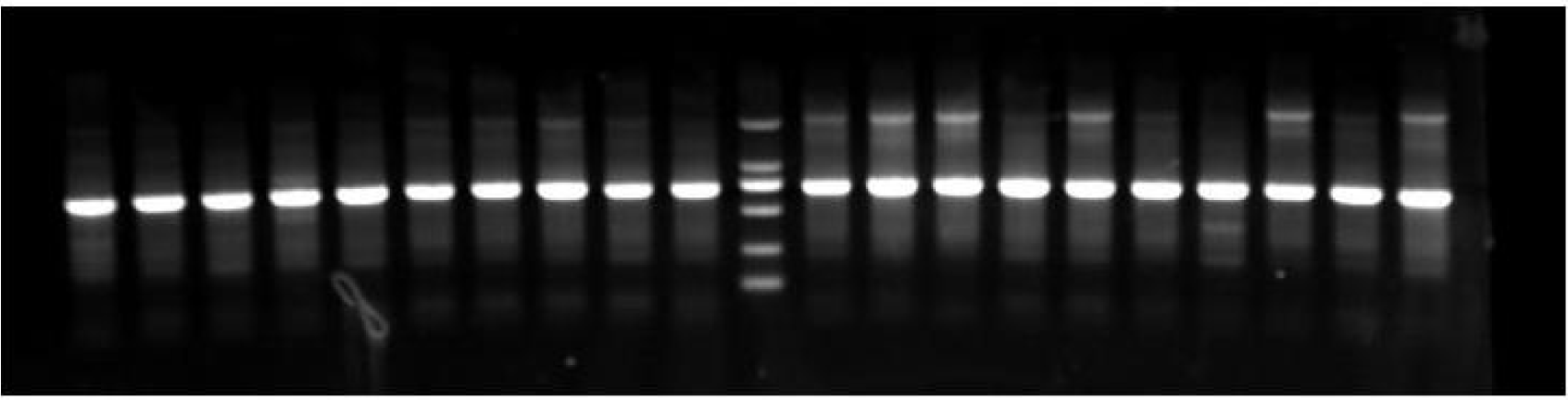
Amplification pattern of primer 12-1

**Fig. 18.**
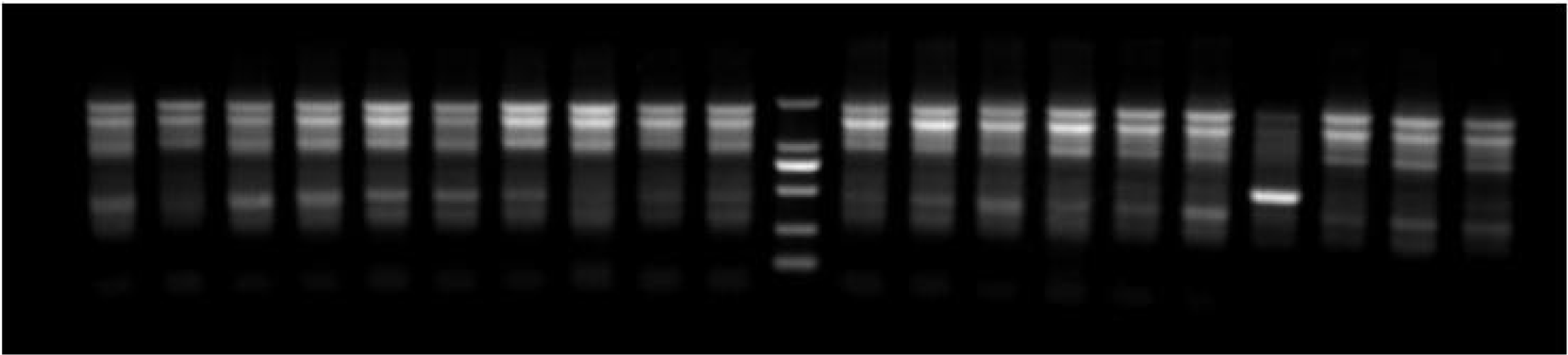
Amplification pattern of primer 12-2

These five electrophoretic maps amplified by primer 9, 10,11,12-1,12-2 show the results of amplification of sequences upstream of six Cao-sequences from the 9th, 10th, 11th, and 12th chromosomes and 10 control individuals on the right. We found that all amplification patterns changed in the 10 mutants.

To obtain in-depth knowledge of the changes occurring in the genome of mutants, whole-genome resequencing was carried out in six selected mutants. A total of 334.05 Gb of data was obtained and Q30 reached over 92.37%. The efficiency of comparison with the reference genome was at least 97.98%, with an average coverage depth of 30×–71× and genomic coverage of 98.45% (at least one base coverage). We prioritized analysis of 5,000-bp-long sequences located upstream and downstream of MBSs, and searched for SNPs, indels, SVs (Structure Variation), and so on.

**Table 4.** Statistics of SNP genotypic sites differing between samples sample

| sample | T1 | T2 | T3 | T4 | ww | ZK |
| --- | --- | --- | --- | --- | --- | --- |
| T1 | 0 | 7216 | 8150 | 6882 | 7691 | 7741 |
| T2 | 7216 | 0 | 7658 | 3725 | 3451 | 6539 |
| T3 | 8150 | 7658 | 0 | 7838 | 7431 | 6873 |
| T4 | 6882 | 3725 | 7838 | 0 | 3433 | 6755 |
| ww | 7691 | 3451 | 7431 | 3433 | 0 | 6767 |
| ZK | 7741 | 6539 | 6873 | 6755 | 6767 | 0 |

**Table 5.** Statistics of the same SNP genotypic loci between samples sample

| sample | T1 | T2 | T3 | T4 | ww | ZK |
| --- | --- | --- | --- | --- | --- | --- |
| T1 | 0 | 6261 | 5327 | 6611 | 5785 | 5782 |
| T2 | 6261 | 0 | 5724 | 9730 | 10002 | 6889 |
| T3 | 5327 | 5724 | 0 | 5560 | 5987 | 6571 |
| T4 | 6611 | 9730 | 5560 | 0 | 10032 | 6694 |
| ww | 5785 | 10002 | 5987 | 10032 | 0 | 6676 |
| ZK | 5782 | 6889 | 6571 | 6694 | 6676 | 0 |

There were 13,867 SNP loci in the six mutants. As indicated in Tables 4 and 5, we found that small differences in SNP loci occurred among the samples T2, T4, and WW, while greater differences in the SNP loci were found among the other samples.

**Table 6.** Statistics of the same indel genotypic loci between samples

| sample | T1 | T2 | T3 | T4 | ww | ZK |
| --- | --- | --- | --- | --- | --- | --- |
| T1 | 0 | 1348 | 1484 | 1273 | 1432 | 1438 |
| T2 | 1348 | 0 | 1403 | 711 | 647 | 1219 |
| T3 | 1484 | 1403 | 0 | 1399 | 1341 | 1224 |
| T4 | 1273 | 711 | 1399 | 0 | 665 | 1228 |
| ww | 1432 | 647 | 1341 | 665 | 0 | 1230 |
| ZK | 1438 | 1219 | 1224 | 1228 | 1230 | 0 |

**Table 7.**
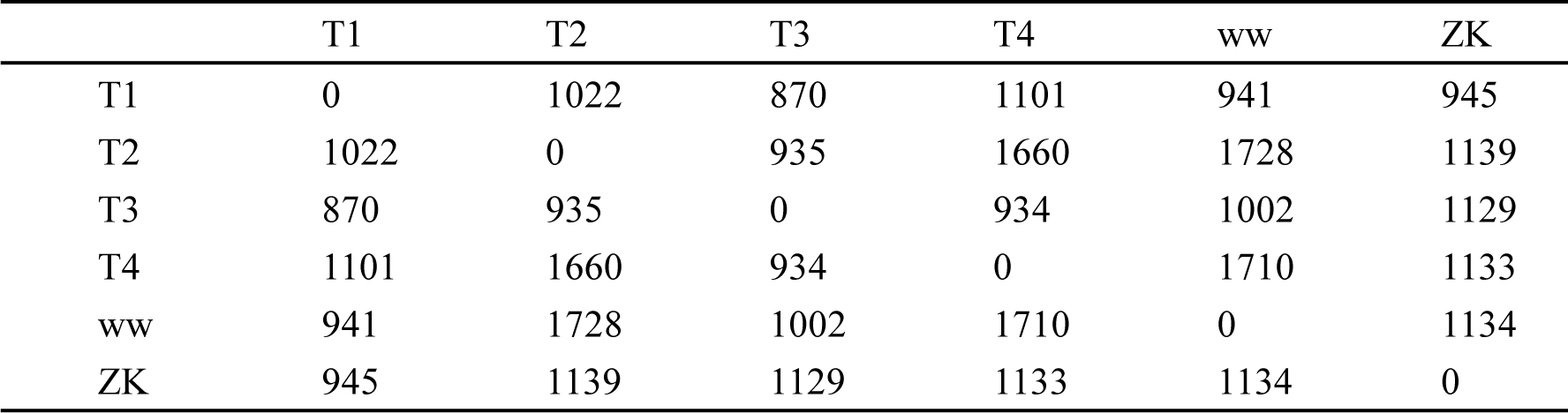
Statistics of the same indel genotypic loci between samples

There were 2,482 indel loci in the six mutants. As indicated in Tables 6 and 7, we found that small differences occurred among the samples T2, T4, and WW in indel loci, while greater differences in the indel loci were found among the other samples.

**Table 8.** Statistics of SV genotype in six mutants

| ID | SV | INS | DEL | INV | ITX | CTX | UN |
| --- | --- | --- | --- | --- | --- | --- | --- |
| T1 | 144 | 3 | 140 | 1 | 0 | 0 | 0 |
| T2 | 136 | 1 | 135 | 0 | 0 | 0 | 0 |
| T3 | 163 | 0 | 161 | 2 | 0 | 0 | 0 |
| T4 | 151 | 2 | 149 | 0 | 0 | 0 | 0 |
| ww | 148 | 1 | 146 | 1 | 0 | 0 | 0 |
| ZK | 170 | 0 | 170 | 0 | 0 | 0 | 0 |

Table 8 indicates that DEL-type mutations accounted for most of the SV mutations.

**Table 9.** Features of deletions in MBS regions

| Chromosome | Deletion of short fragment | Deletion of long fragment | Deletion of super long fragment |
| --- | --- | --- | --- |
| 1 | 5 | 12\18 | 0 |
| 2 | 4 | 13\20 | 1 |
| 3 | 3 | 8\14 | 0 |
| 4 | 1 | 3\17 | 0 |
| 5 | 3 | 10\13 | 0 |
| 6 | 2 | 9\15 | 1 |
| 7 | 6 | 15\21 | 1 |
| 8 | 3 | 9\16 | 0 |
| 9 | 3 | 6\11 | 0 |
| 10 | 0 | 4\7 | 0 |
| 11 | 0 | 6\8 | 2 |
| 12 | 5 | 7\15 | 1 |
| 13 | 1 | 6\9 | 0 |
| 14 | 2 | 8\12 | 0 |
| 15 | 2 | 3\7 | 0 |
| 16 | 3 | 12\20 | 0 |
| 17 | 2 | 7\12 | 1 |
| 18 | 2 | 9\10 | 0 |
| 19 | 1 | 5\10 | 0 |
| 20 | 2 | 10\16 | 0 |
| 21 | 3 | 6\13 | 0 |
| 22 | 0 | 5\7 | 0 |
| 23 | 1 | 7\10 | 0 |
| 24 | 1 | 12\14 | 0 |
| 25 | 0 | 5\7 | 0 |

“Deletion of short fragment” refers to the deletion of a fragment of less than 1,000 bp in length, being mainly distributed upstream of the Cao-sequences. In contrast, “deletion of long fragment” refers to the deletion of a fragment of 1,000 to 10,000 bp in length, which all start at the starting point of the Cao-sequences. Most deletions of long fragments are about 6,100 bp in length; that is, most of the Cao-sequence was deleted, except for about 400 bp left of the tail. The rest involve deletions of Cao-sequences of varying lengths or extending to the downstream sequence. “Deletion of super-long fragment” refers to the deletion of a fragment of more than 10,000 bp, which also started at the beginning of Cao-sequences, ranging up to more than 90,000 bp. A total of 297 Cao-sequence loci were deleted from the six mutants, accounting for 92.2% of the total number of Cao-sequences.

**Table 10.** The common deletion sites of the six mutants

| Chromosome | ratio | location |
| --- | --- | --- |
| 25 | 1\7 | 26600542 |
| 24 | 6\14 | 10539561,15388649,17825677,18261962,19097907,19289796 |
| 23 | 0\10 |  |
| 22 | 1\7 | 31321264 |
| 21 | 0\13 |  |
| 20 | 2\16 | 10259682,21242557 |
| 19 | 2\10 | 24436276,29312438 |
| 18 | 0\10 |  |
| 17 | 3\12 | 32306144,**33880987**,39098463 |
| 16 | 3\20 | 18686244,46064610,**7567349** |
| 15 | 0\7 |  |
| 14 | 1\12 | 48171329 |
| 13 | 1\9 | 12100389 |
| 12 | 1\15 | 31760222 |
| 11 | 3\8 | 14654868,33468804,35114948 |
| 10 | 0\7 |  |
| 9 | 1\11 | 12353529 |
| 8 | 2\16 | 15452959,23450422 |
| 7 | 2\21 | **19787594,48632070** |
| 6 | 7\15 | 22238173,31943883,**35353426**,35361530,41349775,**45762450**,46134553 |
| 5 | 3\13 | **21012267**,32077761,36628049 |
| 4 | 1\17 | 13371759 |
| 3 | 4\14 | **17612411**,19420613,19437684,39658395 |
| 2 | 4\20 | 18401663,20195208,20580757,26570550,**40344073** |
| 1 | 3\18 | 10626533,31037271,48596952 |

The number bold in Table 10 were deletions of short fragments. The rest were all deletions of long fragment except one super-long fragment, which all started at the starting point of Cao-sequences. In the six mutants, 43 of the same locations of Cao-sequences were deleted, accounting for 13.4% of the total. Table 10 shows that there were no common deletion loci in the 10th, 15th, 21st, and 23rd chromosomes, and nearly half of the Cao-sequence loci were common deletion sites in the 6th and 24th chromosomes.

**Table 11.** the to ten result of GO enrichment

| GO.ID | Term | Annotated | Significant | Expected | KS |
| --- | --- | --- | --- | --- | --- |
| GO:0043565 | sequence-specific DNA binding | 687 | 6 | 5.63 | 8.3e-19 |
| GO:0003700 | sequence-specific DNA binding<br>transcriptor | 695 | 6 | 5.7 | 3.4e-10 |
| GO:0003682 | chromatin binding | 198 | 4 | 1.62 | 1.9e-08 |
| GO:0008270 | zinc ion binding | 1342 | 8 | 11 | 2.1e-08 |
| GO:0003924 | GTPase activity | 270 | 3 | 2.21 | 4.1e-08 |
| GO:0019904 | protein domain specific binding | 127 | 0 | 1.04 | 1.3e-06 |
| GO:0016616 | oxidoreductase activity | 113 | 1 | 0.93 | 1.4e-06 |
| GO:0001594 | trace-amine receptor activity | 101 | 0 | 0.83 | 4.0e-06 |
| GO:0005160 | transforming growth factor beta receptor | 40 | 0 | 0.33 | 4.7e-06 |
| GO:0003743 | translation initiation factor<br>activity | 81 | 1 | 0.66 | 5.5e-06 |

GO ID: ID of GO term; Term: function of GO; Annotated: the number of genes from total gene annotated to this function; Significant: the number of genes differentially expressed annotated to this function; Expected: expected value of the number of differential genes annotated to this function; KS: Significant statistics for enrichment of term.

To know what genes were affected after Cao-sequence deletion, the transcriptome sequencing was carried out in four flesh red mutants and three contrast individuals of wild type. Because several different mutants were also found among flesh red mutants, we think that more than epidermis tissues were involved. So we chose the tail part including muscle tissue, epidermis and caudal fin as target tissues. Tthe result of transcriptome sequencing was a comprehensive result of three tissues.

Overall, 702 unigenes were grouped into 25 functional categories using the COG database classification. The dominant groups were ‘general function prediction only’, ‘lipid transport and metabolism’, and ‘signal transduction mechanisn’, ‘energy production andconversion’. A transcriptome comparison revealed that 393 up-regulated genes and 309 down-regulated genes in mutant. These unigenes were annotated to at least one GO term annotation, using the GO classification system. Based on GO annotation, the unigenes were classified into 55 different groups belonging to three main categories: biological process, cellular component, and molecular function. In the biological process category, multi-organism process, rhythmic process, biological phase and cell killing had the obvious proportion difference between contrast and mutant. In the cellular component category, virion, virion part, and nucleoid were the processes with the highest enrichment level difference between contrast and mutant. In the molecular function category, molecular carrier and cargo receptor were significantly different. TopGO enrichment analysis shown the most significant 10 nodes were sequence-specific DNA binding, sequence-specific DNA binding transcriptor, chromatin binding, zinc ion binding, GTPase activity, protein domain specific binding, oxidoreductase activity, trace-amine receptor activity, transforming growth factor beta receptor, translation initiation factor activity. Most were regulation proteins or related ones especially the four top nodes were all DNA bind protein.

**Fig 19.**
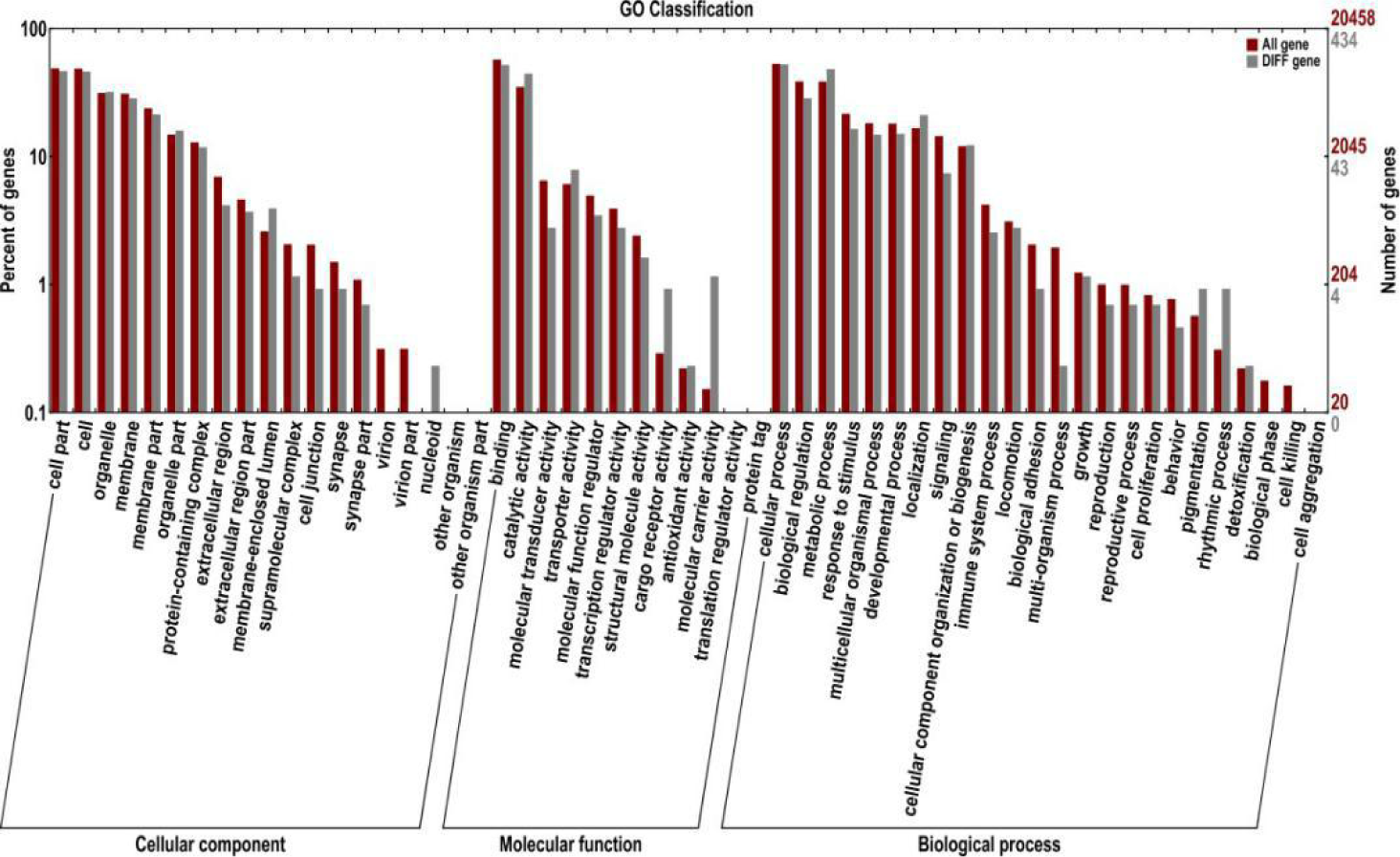
classification chart of differential expression gene with GO annotation

The abscissa is classified as GO, the left side of the ordinate is the percentage of the number of genes, and the right side is the number of genes.The secondary function with obvious proportion difference indicates that the enrichment trend of differentially expressed genes is different from that of all genes.

**Table 12.** the resultof KEGG enrichment of differently expressed genes

| Pathway | KO | Enrichment_Factor | Q_value |
| --- | --- | --- | --- |
| Metabolic pathways | ko01100 | 2.36 | 0.0000e+00 |
| Oxidative phosphorylation | ko00190 | 5.29 | 1.1571e-07 |
| Citrate cycle (TCA cycle) | ko00020 | 10.83 | 9.3627e-08 |
| 2-Oxocarboxylic acid metabolism | ko01210 | 15.81 | 3.2919e-06 |
| Valine, leucine and isoleucine degradation | ko00280 | 6.62 | 4.5650e-05 |
| Fatty acid metabolism | ko01212 | 6.29 | 6.1464e-05 |
| Carbon metabolism | ko01200 | 5.26 | 2.7734e-04 |
| Steroid hormone biosynthesis | ko00140 | 7.91 | 3.1332e-04 |
| Biosynthesis of unsaturated fatty acids | ko01040 | 8.86 | 5.5737e-04 |

Pathway: three level metabolic pathway name of KEGG; KO: ID of KEGG pathway; Q-value: significant statistics of enrichment, the smaller the Q-value, the higher the enrichment degree.

We then performed a KEGG enrichment analysis of the DEGs obtained from tail part of wild type contrast compare with those from mutant. In mutant transcriptomes, the significantly-changed pathways included metabolic pathways, oxidative phosphorylation, citrate cycle, 2-oxocarboxylic acid metabolism, valine, leucine and isoleucine degradation, fatty acid metabolism, carbon metabolism and so on. Most of these pathways were related to basal anabolism, catabolism and energy metabolism.

## Discussion

In 2013, we discovered that introducing rearranged genomic DNA fragments from common carp into fertilized eggs resulted in mutants being produced (Cao et al., 2013a). We subsequently joined the adaptors with rearranged fragments to trace the introduced DNA fragments (Cao et al., 2013b). Because few of the introduced DNA fragments integrated into the genome, it was very difficult to obtain these fragments from the genome by one-step amplification. Therefore, we designed longer adaptors and two primers for a two-step amplification process based on the adaptor sequences. The first amplification product, whose template was the second digestion product with adaptors, was microinjected into fertilized zebrafish eggs. This step ensured that most of the microinjected fragments carried the adaptor. We were then able to easily introduce sequences from the two-step amplification, and the second amplification product was directly microinjected into fertilized zebrafish eggs. The successful application in zebrafish demonstrated that introducing rearranged genomic DNA into a host can produce mutant characters.

This study found that some of the endogenous DNA fragments that had been rearranged in sequence were mutagenic.This study found that a sequence has mutagenicity, which exhibited two unique features. The first was that its integration site was highly fixed; all such sites were upstream of a long 6,537-bp C-sequence, which has neither coding function nor similarity to any regulatory structure. This has never been reported in previous studies. The second feature was that, after introduction of the sequence into fertilized zebrafish eggs, it could change into another highly similar sequence. The changes happened in the sequences amplified from mutants include that the original enzyme cutting sites disappeared, and there was a highly variable region found among these sequences, nearly every individual differed from each other in the region. Therefore, we think that the introduced sequence is reformed before integration into the genome, although the specific process involved in this needs further research.

The result of whole-genome resequencing showed that the Cao-sequences were deleted in different locations after introducing the sequences into fertilized zebrafish eggs, and the deletion positions of different individuals were not identical. Upon combining all experimental results, we speculated that, after the introduction of the sequence into the fertilized egg, it becomes linked to Cao-sequence by an unknown mechanism. Subsequently, the introduced sequences pair and recombine with the homologous sequence, followed by the introduction of Cao-sequence into a new location while part of it (mostly 6,100 bp from the start of the Cao-sequence) is cut off, leaving a partial Cao-sequence remnant in situ. The introduced sequences that were the same in highly variable region preferentially associated with the Cao-sequence at certain locations. The change of highly variable regions could modify this preference, causing a wider deletion of Cao-sequence after introduction of the mixed sequences, resulting in the death of the host.

The result of transcriptome analysessuggested most of the genes affected in the mutants were regulatory factors and DNA-binding proteins that regulates gene-expression, as well as some receptor genes. KEGG enrichment analysis results shown that differentially expressed genes were mainly concentrated in some basal metabolic pathway. We speculated that those genes encoding regulatory proteins (such as transcription factors, DNA binding proteins, receptors, etc.) in the zebrafish genome were divided into 322 regulatory units. Each regulatory unit control the expression of a number of functional genes through these regulatory proteins and form a basic physiological function. It is similar to large computer programming and we call these regulatory units genetic modules. Cao-sequence may be the module boot sequence and excessive deletion of Cao-sequence or inappropriate Cao-sequence combinations deletion can lead to the loss of many important underlying functions and lead to death. In future work, we need to analyze the basic functions of these 322 modules through a large number of mutants combined with resequencing. It would be interesting to determine whether the character of zebrafish is similar to the design of a network APP, which is directly formed by different modular combinations.

The technique presented in this paper originated from the results of research on crops and was also shown to be effective in the study of common carp, so we speculated that the results are likely to be applicable to most plants and other animals, including humans. However, in a search of NCBI, we did not find any homologous sequences similar to Cao-sequence in other species. Therefore, we speculated that each species may have its own unique guidance sequences.

## Materials and Methods

### 1. Extraction of the zebrafish genome

Eighty zebrafish embryos were collected 5 days after fertilization and incubated with 500 μl of lysis solution containing 5 μl of protease K at 55° for 4 h. After lysis, 3 μl of RNase A (10 mg/ml) was added and maintained at 37° for 30 min. A 500-μl mixture of phenol and chloroform (1:1) was added, vortex-mixed for 10 s, centrifuged at 12,000 × g for 10 min, and then 300 μl of the supernatant was transferred to another centrifuge tube. A 2.5-fold volume of ethanol was added, the mixture was centrifuged at 12,000 × g for 10 min after mixing, the supernatant was discarded, and then 1 ml of 75% ethanol was added and mixed. Next, the solution was centrifuged again at 12,000 × g for 10 min and the supernatant was discarded. The precipitate was dried at room temperature and dissolved in 50 μl of ultrapure water at 50°.

### 2. Restriction digestion and rearrangement of zebrafish genomic DNA

Zebrafish genomic DNA (10 μg) was digested in a 200-μl volume with 10 μl of *Msp I* and 20 μl of digestion buffer at 37°C for 4 h. A 4-μl aliquot of the digested products was subjected to 0.8% agarose electrophoresis. The remaining 196 μl of digested products was placed in a water bath at 95°C for 10 min to inactivate the *Msp I*. Then, 104 μl of ultrapure water and a 300-μl mixture of phenol and chloroform (1:1) was added, vortexed-mixed for 10 s, and centrifuged at 12,000 × g for 10 min. A 250-μl aliquot of the supernatant was transferred to another centrifuge tube. Twenty-five microliters of 3 M NaoAC (pH 5.4) and a 2.5-fold volume of ethanol were added for precipitation (−80°C for 2 h), and the solution was then centrifuged at 12,000 × g for 10 min. The supernatant was discarded, after which the precipitate was washed twice with 70% ethanol and then dried at room temperature. The precipitate was next resuspended in 45 μl of ultrapure water and directly added to T4 DNA ligase (4°C overnight). The ligation reaction system included 60 μl (45 μl of restriction fragment, 3 μl of T4 DNA ligase, and 12 μl of 5× T4 DNA ligase buffer). One microliter of the rejoined product was subjected to 1% agarose gel electrophoresis and the remaining 59 μl was incubated at 70°C for 10 min to inactivate the T4 DNA ligase. Then, 241 μl of ultrapure water was added to a 300-μl mixture of phenol and chloroform (1:1), vortex-mixed for 10 s, centrifuged at 12,000 × g for 10 min, and a 250-μl aliquot of the supernatant was transferred to a new centrifuge tube. Twenty-five microliters of NaoAC (3 M pH 5.4) and 2.5 volumes of ethanol were then added for precipitation (−80°C for 2 h), and the mixture was centrifuged at 12,000 × g for 10 min. After removing the supernatant, the precipitate was washed twice with 70% ethanol and dried at room temperature. Finally, the precipitate was resuspended in 55 μl of ultrapure water.

### 3. Restriction digestion of the rearranged genomic DNA

Fifty-two microliters of rearranged DNA was incompletely digested with the EcoR I enzyme in a 20-μl reaction system, including 1 μl of EcoRI, 2 μl of 10× digestion buffer, and 17 μl of rearranged genomic DNA, for 20–40 min. The DNA fragments were recovered from the dispersion zone corresponding to fragment lengths greater than 2 kb using the Axygen Gel Extraction Kit (Axygen Scientific Inc., Union City, CA, USA). The product was resuspended in 20 μl of ultrapure water.

### 4. Adding the adaptor

The DNA fragments purified with the agarose gel DNA purification kit were mixed with Gmadaptor1 and Gmadaptor2 (100 μM, Gmadaptor1:

GTCATCTCAAACCATCTACACGGAACCAAACACATGAAGCCG; Gmadaptor2: AATTCGGCTTCATGTGTTTGGTTCCGTGTAGATGGTTTGAGATGA) (20-μl reaction volume, including 2 μl of each primer, 4 μl of 5× T4 DNA ligase buffer, and 12 μl of purified DNA fragments), heated to 95°C for 5 min, cooled, and then bound directly with T4 DNA ligase (20-μl reaction volume, 1 μl of T4 DNA ligase, 4 μl of 5 ×T4 DNA ligase buffer, and 15 μl of the above products) at room temperature for 2 h. The product was purified using a DNA fragment purification kit. The recovery solvent was 20 μl of ultrapure water.

### 5. PCR amplification

Amplification was performed with Gmprimer1 (5 μM, TCATCTCAAACCATCTACACG). The reaction volume was 50 μl, including 25 μl of 2× Mastermix (Vazyme Biotech Co., Ltd., Nanjing, Jiangsu, China), 5 μl of primer, 18 μl of ultrapure water, and 2 μl of the purified recovered DNA fragments. The following amplification protocol was followed: pre-denaturation at 95° for 2 min; 33 cycles of denaturation at 95° for 30 s, annealing at 50° for 45 s, and extension at 72° for 2 min; followed by final extension at 72° for 5 min. A 2-μl aliquot of the amplification products was subjected to 1% agarose gel electrophoresis. The remaining amplification products were recovered with the Axygen PCR clean up kit, and the recovery solvent was 20 μl of ultrapure water. One microliter of the recovered product was measured in a UV spectrophotometer.

### 6. First microinjection

The recovered amplified product was diluted with ultrapure water to a working concentration of 25 ng/L, which was used to microinject fertilized zebrafish eggs. The microinjected volume per embryo was about 1 nl. A blank contrast (injection of a 2-kb product that was unrelated to the amplified product) was prepared and injected into control embryos. The embryos were reared in incubators at 28.5°C.

### 7. Genomic DNA extraction from mutant and amplification of the introduced sequence

After a 3-month breeding cycle, we collected the individuals that exhibited clear phenotypic differences from normal zebrafish for further study. The genomic DNA extraction methods were the same as those used in the first amplification step. The first amplification was performed with Gmprimer1. The reaction volume was 50 μl, including 25 μl of 2× Mastermix (Vazyme Biotech), 5 μl of primer, 18 μl of ultrapure water, and 2 μl of the mutant genomic DNA. The amplification protocol was: pre-denaturation at 95° for 2 min; 33 cycles of denaturation at 95° for 30 s, annealing at 50° for 45 s, and extension at 72° for 2 min; followed by final extension at 72° for 5 min.

The second amplification was performed with Gmprimer2 (5 μM, CCAAACACATGAAGCCGAATT). The reaction volume was 50 μl, including 25 μl of 2× Mastermix (Vazyme Biotech), 5 μl of the primer, 18 μl of ultrapure water, and 2 μl of the PCR products from the first amplification. The amplification protocol was: pre-denaturation at 95° for 2 min; 33 cycles of denaturation at 95° for 30 s, annealing at 50° for 45 s, and extension at 72° for 2 min; followed by final extension at 72° for 5 min. A 2-μl aliquot of the second-step amplification products was subjected to 1% agarose gel electrophoresis. The remaining amplification products were recovered with an Axygen PCR clean up kit. The recovery solvent was 20 μl of ultrapure water. The largest amplified band from the mutant was collected to be cloned and sequenced.

### 8. Second microinjection

The recovered second amplified product was diluted with ultrapure water to a working concentration of 25 ng/L, which was used to microinject fertilized zebrafish eggs. The second microinjection method followed the seven steps described above, and the surviving embryos were raised until an age of 3 months to identify the mutants.

### 9. Genomic DNA extraction of mutants after the second microinjection and amplification of the introduced sequence

The method to extract genomic DNA from the mutant after the second microinjection was the same as that described above. The introduced sequence was amplified directly with the primer Gmprimer2 and the process followed the eight steps described above using the template from extracted genomic DNA of the mutant. The amplified sequences were cloned and sequenced.

### 10. Third microinjection

Ten recovered amplified products from 10 mutants were mixed equally for the third microinjection as described above. This mixture was diluted with ultrapure water to four working concentrations of 25, 5, 0.5, and 0.25 ng/L, which were used to microinject fertilized zebrafish eggs. The microinjected volume per embryo was about 1 nl. A blank contrast (injection of a 2-kb product that was unrelated to the amplified product) was prepared and injected into control embryos. The embryos were reared in incubators at 28.5°C.

### 11. Genome walking

The program of genome walking was performed using Genome Walking Kit (No. 6108) produced by Takara Biomedical Technology. The three primers for the downstream sequence were f1: CTGGCATTCGTTCACTCAAGAA; f2: TGAATATGATACAGCTGAATAA; and f3: CTACCTGTCTCTTACATTTC. The three primers for the upstream sequence were: r1: tgtatttaagccaaaacacaac; r2: aaatcaataaattaatacgtggg; and r3: tgtatacaaacctttcttgtcg.

### 12. Amplification of upstream sequence of MBS

To detect the changes in the upstream sequence of MBS, we designed two primers specific to regions within MBS: R2, attccgtcgccataatgact; and R1, taaattagggccagggacac. At the same time, we designed six primers specific for regions located in the sequence upstream of MBS with different locations in the 8th chromosome, along with five primers in the sequence upstream of MBS from the 9th, 10th, 11th, and 12th chromosomes.

**Table 1.**
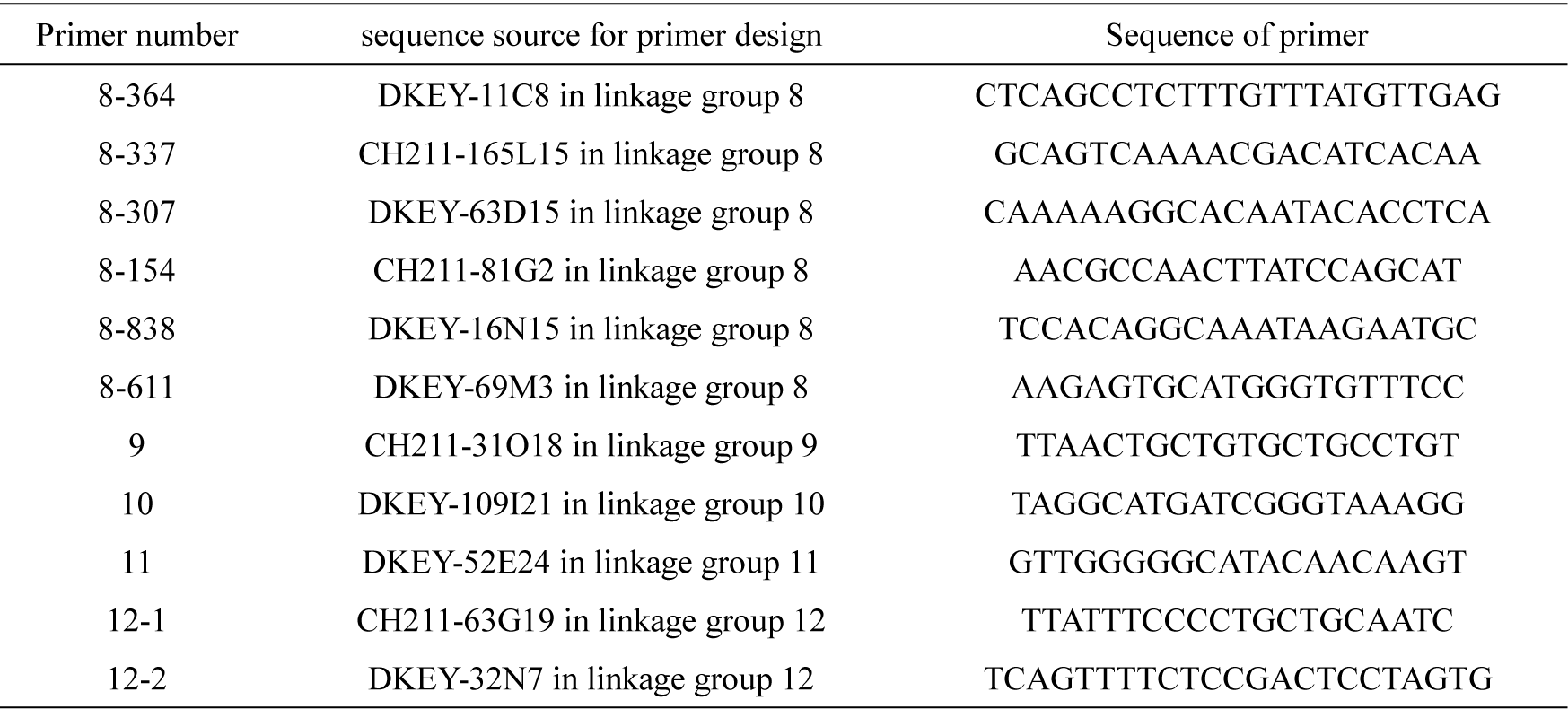
Primers for analysis of region upstream of MBS

### 11. Whole-genome resequencing of zebrafish mutants

Whole-genome resequencing of zebrafish mutants was carried out by Genepioneer Biotechnologies (reference genome version: Danio_rerio.GRCz10.dna.toplevel.fa; URL: ftp://ftp.ensembl.org/pub/release-91/fasta/danio_rerio/dna/Danio_rerio.GRCz10.dna.toplevel.fa).

**Table 2.** number of six samples and description of mutant characters

| Number of sample | Character description |
| --- | --- |
| T1 | red flesh color |
| T2 | red flesh color |
| T3 | red flesh color |
| T4 | red flesh color |
| ww | curved stripe |
| ZK | the mouth can't close |

### 12. Transcriptome sequencing of zebrafish mutant

The muscle tissues from four red flesh mutants and three wild type individuals were collected for transcriptome sequencing. Transcriptome sequencing and data analysis of zebrafish mutants and contrast was carried out by Genepioneer Biotechnologies.

## Acknowledgments

We thank Liwen Bianji, Edanz Group China (www.liwenbianji.cn/ac), for editing the English text of a draft of this manuscript.

